# Fusion of lipid membranes: an alternative pathway

**DOI:** 10.64898/2026.02.19.706911

**Authors:** Di Jin, Zhi Li, Sylvie Roke, Jacob Klein

## Abstract

Membrane fusion is ubiquitous in myriad biological processes, including presynaptic neurotransmitter release, myoblast fusion, virus entry, and fertilization^1–3^. It is often considered in the presence of fusogenic proteins, which mediate the fusion by pulling the membranes into contact to overcome the high energy barrier imposed by hydration repulsion^4–7^. Here we uncover an alternative pathway – membrane fusion arising from electroporation induced by transmembrane potentials^8,9^ in absence of proteins. Using biased molecular dynamics simulations to carry out extensive potential-of-mean-force calculations^10^, we show how electroporation of opposing membranes promotes membrane fusion via formation of an intermembrane splayed lipid and peripore stalk, without any protein involvement. We validate our mechanism by demonstrating experimentally, via video microscopy and second harmonic generation imaging^11^ of interacting giant unilamellar vesicles (GUVs), that GUV fusion occurs in the presence of transmembrane potentials but not in their absence. The magnitude of these transmembrane potentials is biologically relevant, prevalent in transient states near the surfaces of cell membranes^8,12^, underscoring the potential importance of this mechanism in biological events.

## Main

Cell-cell fusion is largely controlled by the structural properties of the biological membranes ^1–3^, a reductionist model of which consists of a lipid bilayer. The lipidic headgroups forming the membranes’ surface are highly hydrophilic and hydrated, imposing a high energy barrier for two membranes to fuse due to hydration repulsion^4–7^. Fusogenic proteins—most notably SNAREs—are widely recognized as key drivers of membrane fusion because by pulling the membranes together to under 1 nm they reduce lipid hydration and thus lower the hydration-repulsion energy barrier ^7,13^.

This greatly increase the likelihood that lipids adopt the *splayed* state, in which one acyl chain inserts into the opposing membrane—an energetically costly early fusion intermediate^14^. Such intermembrane splayed lipids are thought to template the reorientation of surrounding lipids, promoting formation of the next intermediate, the *stalk*: a nanometric lipid bridge connecting the two proximal monolayers ^14–17^. The stalk is metastable and can relax into a hemifusion diaphragm, where the bilayers merge into a single sheet. Complete fusion then proceeds by formation and expansion of a fusion pore within this diaphragm, yielding a fully fused membrane^5,7^, as illustrated in Fig. 1A.

**Fig. 1.**
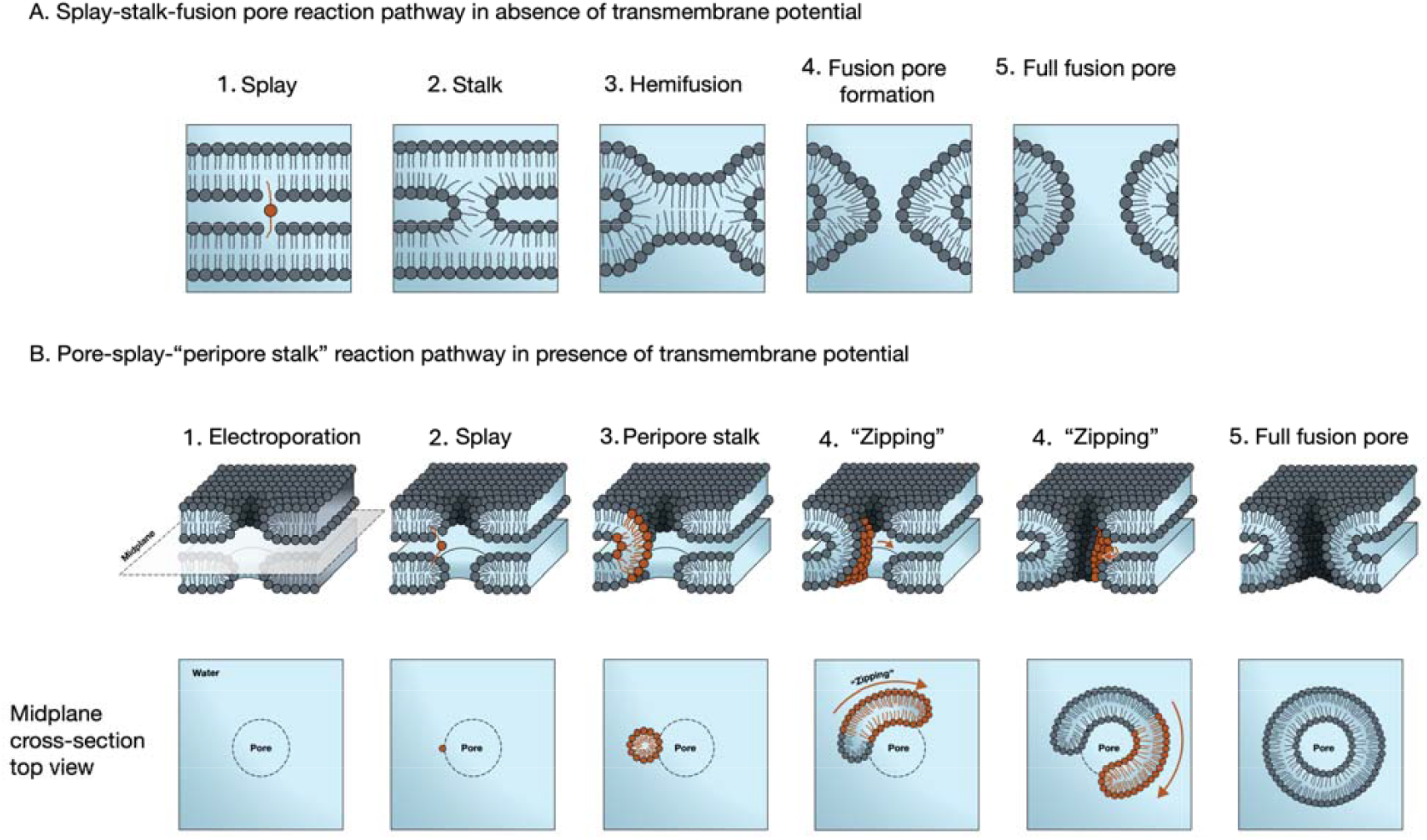
Schematic illustration of two distinct membrane fusion pathways occurring at the point of contact between two lipid bilayers. (A) In the absence of transmembrane potential, fusion proceeds via the classical pathway, where sufficiently close approach of the membranes (facilitated, for example, by fusogenic proteins) results in lipid splaying (1), and stalk formation (2), followed by hemifusion (3), and the nucleation of a fusion pore (4, 5). (B) In the presence of a transmembrane potential, fusion may proceed via a different pathway. Here, electropores form first (1), promoting splayed lipid (2), followed by peripore stalk formation (3), by disrupting the membrane interface, with no need for dehydration by specialized proteins. An annular “zipping” process (4), initiated by the peripore stalk, then zips these pores together, driven by the lower line tension of the pore edges, sealing the membrane contact, to the final fusion pore as shown (5).

Considerable understanding of these intermediates has emerged over the past decades, aided by molecular dynamics simulations based on lipid-only systems^13,14,18^. A consistent conclusion is that fusogenic proteins are indispensable because they facilitate the otherwise unfavorable intermediates of splayed lipids, stalks, and fusion pores. Yet the main mechanism of fusion remains the lipid-mediated topological rearrangements, with proteins largely acting to promote the membrane configurations required for these steps^19,20^ (Fig. 1 A(1)).

Here we describe and validate an alternative physicochemical mechanism for fusion between interacting lipid membranes, which operates independently of fusion proteins, even at the earliest stages, without requiring sub-nanometric membrane contact. Rather, fusion is initiated by the formation of pores^21^ in the opposing membranes stimulated by the electric field. These pores, referred to below as the electropores, give rise to the initial hydrophobic lipidic connection between the membranes, a *peripore stalk*, Fig. 1B(3). This contrasts sharply with the traditional stalk pathway between planar bilayers described above^14–17^, (Fig. 1A). The peripore stalk involves both the proximal and distal monolayers of the two membranes: it is not a hemifusion stalk, as we shall see, since it bridges the whole membranes rather than solely the proximal leaflets (hemi-membranes). Therefore, topologically, the peripore stalk is a direct intermediate of full fusion (Fig. 1B(3–5)), while the traditional stalk is an intermediate of hemi-fusion (Fig. 1A(3)).

Electropores may form under external electric fields^9^ and can also arise spontaneously in naturally-occuring (particularly biological) systems due to transient transmembrane potentials arising from bilayer-surface charge flucuations^8,12,22–30^. These pores not only promote peripore stalk formation but, as we show, persist along the fusion reaction coordinate and transform directly into a fusion pore, bypassing hemifusion. Our results further reveal a “zipping” process, as indicated schematically in Fig. 1B(4), in which a localized peripore stalk expands annuרשאיקרlarly around the rims of opposing electropores to produce the final fusion pore, eliminating the need for a hemifusion diaphragm. This energetically favorable pathway reduces pore-edge line tension and thus proceeds spontaneously^31^. Our analysis employs biased molecular-dynamics simulations building on pioneering work on splayed lipids and fusion stalks^10,13,14,18,32,33^, and is motivated by our earlier finding that the pronounced interfacial dissipation observed between sliding bilayers under transmembrane electric fields arises from spontaneous stalk formation^9^. Finally, we validate our conclusions by demonstrating experimentally, via video and second harmonic generation imaging, that fusion occurs between lipid membranes, in the form of GUVs subject to spontaneous (i.e. not externally applied) transmembrane potentials, with no need for any fusogenic agents; while in the absence of such transmembrane potentials, no GUV fusion occurs. Clarifying this fusion mechanism may advance not only our understanding of cell-cell fusion, but support the development of fusion-dependent technologies such as liposomal drug delivery^34–36^.

### Electropores lower the energy barrier for peripore-stalk formation

Our proposed membrane fusion pathway centers on electropores, which promote metastable inter-bilayer splays, lowering the energy barrier for peripore stalk formation even at membrane separations >1 nm — conditions under which strong hydration repulsion would otherwise hinder stalk formation^4,7,37^. We demonstrate this first by comparing the peripore stalk formation energy barriers ΔG_barrier_ obtained from potential of mean force (PMF) calculations with and without electroporation; at the same time, the splay-formation energetics show that electropores can effectively match the energy-barrier-reducing effect of dehydration. We further confirm this mechanism by observing directly the process of zipping of the electropore rims to form the final full-fusion pore.

Fig. 2 A–C show that unporated bilayers form a fusion stalk, while electroporated bilayers (Fig. 2D-F) form a peripore stalk. The peripore stalk structure shown in Fig. 2E is representative of the morphology observed across a range of midplane separations (Figs. S1-S2). While variations arise from the relative positions of the two electropores, the overall topology is preserved, reflecting the natural translational and rotational freedom of interacting lipid bilayers. Both electroporated and pore-free processes exhibit similar stalk-formation barriers (ΔG_stalk_ ≈ 50 kJ/mol ≈ 20 k_B_T; Fig. 2 G–H), though, notably, at significantly-different bilayer separations *D*_PO4_ across the midplane. In particular, we see clearly that under electroporated conditions (Fig. 2H), more PMF profiles already around *D*_PO4_ ≈ 1 – 1.1□nm show a clear local maximum, indicating metastable stalk formation, while in the absence of electropores (fig. 2G) such local maxima are seen only at D < ca. 0.7 nm. As summarized in Fig. 2I, ΔG_stalk_ at *D*_PO4_ ≈ 0.92 nm, under electroporated condition (Fig. 2D–F) is comparable to that at *D*_PO4_ ≈0.61 nm for the unporated condition (Fig. 2A–C). The latter system corresponds to extreme dehydation of *n*_w_ **=** 5 water/lipid at the midplane, considering that the hydration shell of unperturbed POPC is *n*_w_ **=** 10-12 water/lipid^37–39^, while indeed *D*_PO4_ ≈ 0.9 nm in the electroporated bilayers corresponds to *n*_w_ ≈ 11-13 water/lipid at the midplane (SI Table 1).

**Fig. 2.**
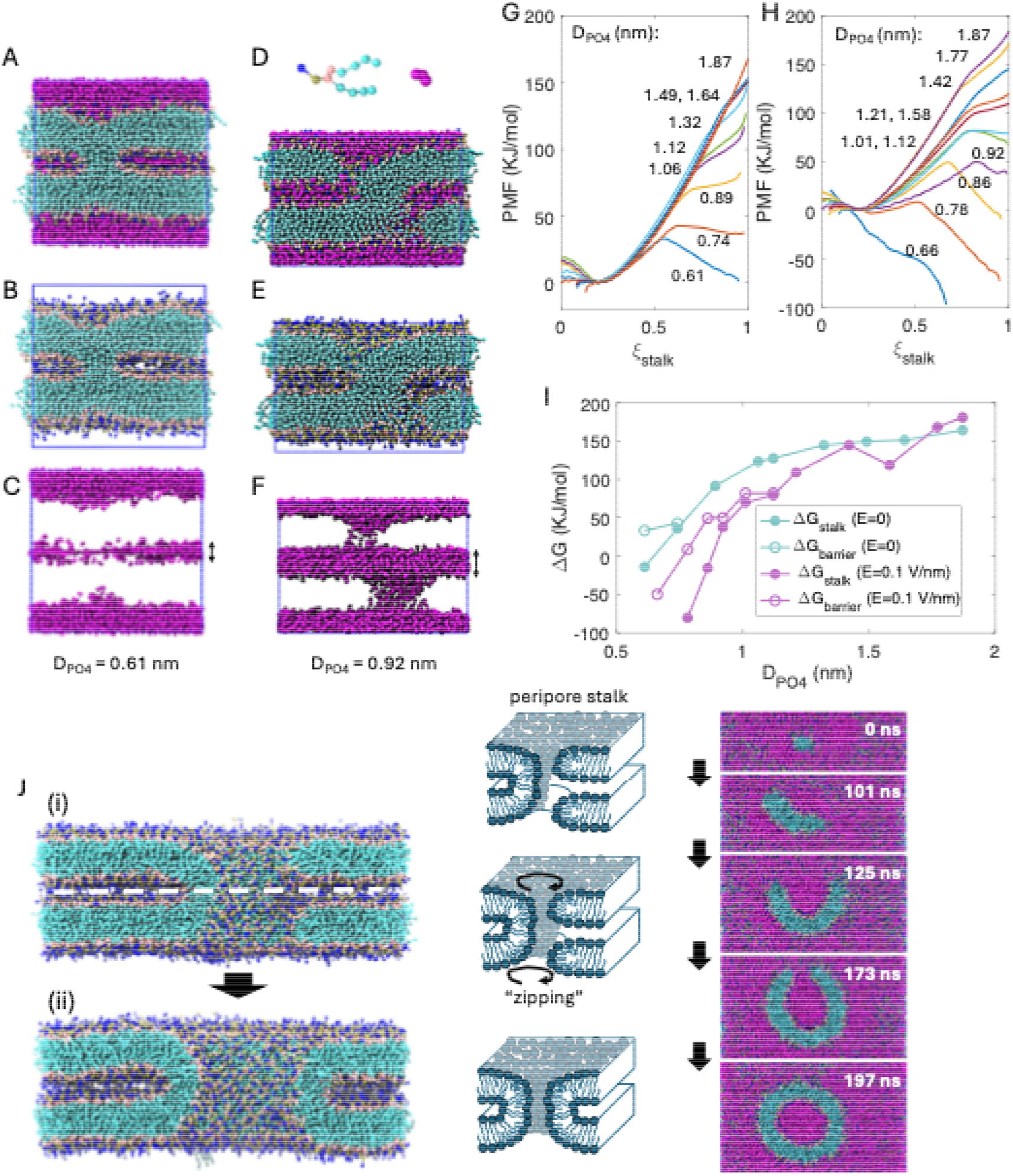
(A–F) Representative simulation snapshots of stalk formation without an electric field (A–C), and peripore stalk formation with an applied electric field (D–F) on Palmitoyloleoylphosphatidylcholine (POPC) bilayers. Periodic boundary conditions are applied across the blue box. Molecular groups color: blue = choline, gold = phosphate, pink = glycerol, cyan = hydrocarbon chain, purple = water. (D, inset) Coarse grained Martini POPC molecule and polarizable water bead. (B,E) Showing only lipids. (C,F) Showing only water. (G) and (H): Potential of mean force (PMF) profiles for systems with varying bilayer separations under *E* = 0 (G) and *E* = 0.1 V/nm (H). The bilayer separation across the midplane, *D*_PO4_,indicated by the arrows in C, F, is the distance between the average *z-*positions of the phosphate groups in opposing leaflets, and is labeled on the curves in G, H. The reaction coordinate, ξ_stalk_, ranges from 0.2 (meta-stable lamellar phase) to 1.0 (fully formed stalk). (I) Energy barriers extracted from (G) and (H). ΔG_stalk_ is the free energy difference at ξ_stalk_ = 0.99 relative to ξ_stalk_ = 0.2, and ΔG_barrier_ corresponds to the local maximum along the reaction coordinate. (J) MD simulation snapshots of the initial peripore-stalk/electropore configuration, fig. J(i)), and final fusion pore (fig. J(ii)) following zipping. On the right, representative snapshots at increasing times from a top view across the midplane section – broken white line in fig. J(i) – showing the zipping process to the final fusion pore.

A possible simple interpretation of the computational results in macroscopic physical terms is that the formation of the peripore stalk is enabled by the high curvature of the lipid monolayer covering the rim of an electropore. The surface of the rim is extremely strongly curved in one direction, with the curvature radius equal to the monolayer thickness. Consequently, this curvature may strongly reduce the energy of the hydration repulsion between the rims of the opposing electropores compared to that between two flat membranes, analogous to the electrostatic repulsion between two strongly curved charged surfaces (e.g., cylinders) versus flat planes of equal charge density. For curved surfaces, hydration repulsion decays faster with distance. At the same time, the large curvature of the electropore rim must drastically decrease the change in elastic energy associated with peripore stalk formation. Because the stalk forms from the opposing electropore rims, its formation is accompanied by relaxation of the bending energy of the rim elements, replaced by the elastic energy of the stalk. This relaxation significantly lowers the overall energy cost compared to regular stalk formation between flat membranes, in which no such relaxation occurs.

Once a peripore stalk forms, we see clearly, in fig. 2J, the time progression, illustrated by the MD simulation snapshots, of how it zips together two opposing overlapping electropores, going from the initial stalk/electropores configuration (fig. 2J(i)) to the final bilayer fusion pore (fig. 2J(ii)), as indicated in the schematic cartoon and the time-series of the zipping process. See also Movie S1.

### Electropores promote and stabilize the formation of the splayed lipid state

What is the origin of the lowered energy barrier for peripore stalk formation between electroporated membranes, that leads to their fusion as seen above? Previous cornerstone studies established that the splayed state formation, as in Fig. 1, and also Fig. 3A, is an essential precursor to stalk formation ^13,14,18^. We examined therefore whether the enhanced formation of peripore stalks could be attributed to more readily formed lipid splays between electroporated bilayers. To address this question, we investigate the energetics of splay formation using umbrella sampling^40^, both with and without electropores. Here the reaction coordinate (ξ) is defined as the distance between the center of mass of the unsaturated tail—known to more readily traverse the bilayer—and the midplane of the opposing leaflet^14^ (Fig. 3B). The resulting PMF profiles typically exhibit two distinct energy barriers, as indicated in Fig. 3C: ΔG□, corresponding to the barrier for initiating splay formation, and ΔG□, associated with the retraction of a splayed lipid back into the lamellar state. A finite ΔG□ indicates a metastable splayed configuration. In each system, a small fraction of splayed lipids lack a finite ΔG□, indicating instability of the splayed state (Fig. S6A).

**Fig. 3.**
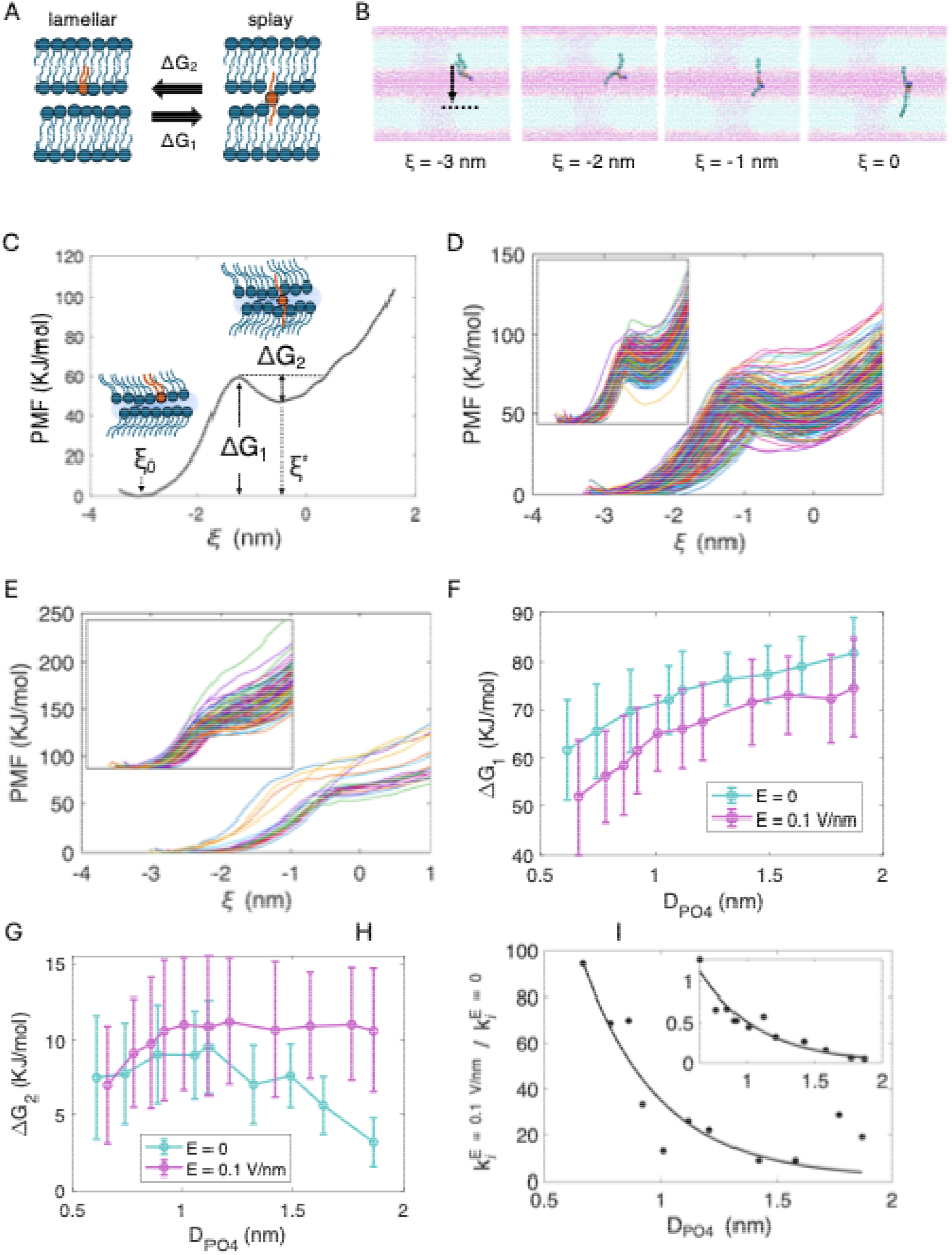
Simulations of splay formation. (A) Single-molecule transition between the lamellar and splayed states, defining the associated energy barriers. (B) Snapshots of splay formation, with the reaction coordinate ξ defined as the distance between the center of mass (COM) of the unsaturated POPC tail and the midplane of the opposing leaflet. (C) Example PMF profile of a single splay event. (D–E) Statistical analysis of energy barriers for the *E* = 0.1 V/nm system with *D*_PO4_ = 0.92 nm, and *E* = 0 system with *D*_PO4_ = 0.61 nm: (D) PMF profiles acquired with the *E* = 0.1 V/nm system which show a clear metastable splay, indicated by a local minimum and a finite ΔG□. (Inset: *E* = 0 system, same axes range). (E) PMF profiles acquired with the *E* = 0.1 V/nm system without a local minimum, indicating unstable splay formation. (Inset: *E* = 0 system, same axes range). (F – G) Energy barriers for splay formation as a function of bilayer midplane separation. Error bars indicate standard deviation. (H) Relative reaction rate of forward splay transitions (lamellar → splay) under *E* = 0.1 V/nm compared to *E* = 0 conditions. Inset: relative reaction rate of backward transitions (splay → lamellar) under *E* = 0.1 V/nm compared to *E* = 0 conditions.

Fig.3D-E present data for the electroporated system (Fig. 2D, *D*_PO_□ = 0.92□nm) and the unporated system (Fig. 2A, *D*_PO□_ = 0.61□nm). The figures show PMFs of metastable splays (Fig. 3D) and unstable splays (Fig. 3E). Notably, the distribution of the splay formation energy barrier ΔG□ (Fig. S7) is comparable in both systems (~20□k_B_T), despite the large disparity in *D*_PO4_, supporting our earlier findings in Fig. 2.

In Fig. 3F–G, we plot ΔG□ and ΔG□ as functions of bilayer midplane separation. It is very clear that electropores lower the energy barrier for splayed state formation (ΔG□) while raising the barrier for the reverse process (ΔG□). Notably, ΔG□ persists at a finite value of 10 KJ/mol (~ 4 k_B_T) even at separations *D*_PO□_ up to 1.9□nm, indicating that splayed lipids formed in the presence of electropores have significantly longer lifetimes. We estimate the reaction rates using an Arrhenius-like formulation described previously. The rate of the reaction can be described as 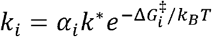 where *i* = splayed state or lamellar state, k^*^is a characteristic relaxation rate of lipids in lamellar state and *α*_*i*_ is the proportionality constant. To evaluate the effects of electroporation, we plot the ratio of the rates: 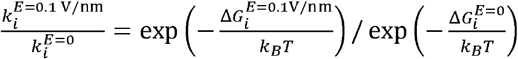. As shown in Fig. 3H, the rate of splay formation increases by an order of magnitude or more under electroporated conditions (e.g., a ca. 25-fold faster splay formation for *D*_PO4_ = 1 nm), while the splay retraction rate is reduced by about half. These results further support the conclusion that electropores both facilitate splay formation and stabilize the splayed state. We attribute this enhanced splayed-lipid formation to the increased topological complexity when pores are present, which broadens the energy landscape. Statistical analysis of over 5000 PMF profiles of splay formation shows that the energy landscape is highly sensitive to a lipid’s vertical position.

Electropores enhance vertical fluctuations and thin the bilayer by sequestering water, expanding the accessible configurational space and enabling splay formation along lower-barrier pathways while stabilizing it at lower free-energy states (Supplementary Analysis).

### Fusion of giant unilamellar vesicles (GUVs) is enabled by spontaneous transmembrane potentials

To validate our mechanism of electropore-facilitated membrane fusion, we examined the interactions of GUVs, which are established models for lipid bilayer membranes^41,42^, emulating cell membranes to some extent. Optical video-monitoring of contacting GUVs^43^, while simultaneously imaging them with second harmonic (SH) generation, enabled us to correlate their interactions with the spontaneous (i.e. not externally applied) potentials ΔΦ_0_ across their membranes^11,44^. As noted, such potentials arise naturally due to bilayer-surface-charge/electric double layer fluctuations, and result in transient membrane electropores^8,12^. As shown in Fig. 4, we find that when transmembrane potentials are present across the bilayers, with no external fusogenic agents or fields, the adjacent GUVs fuse; while in the absence of any spontaneous transmembrane potential, e.g. when membranes are composed of zwitterionic lipids only, no fusion of the GUV membranes occurs.

**Figure. 4.**
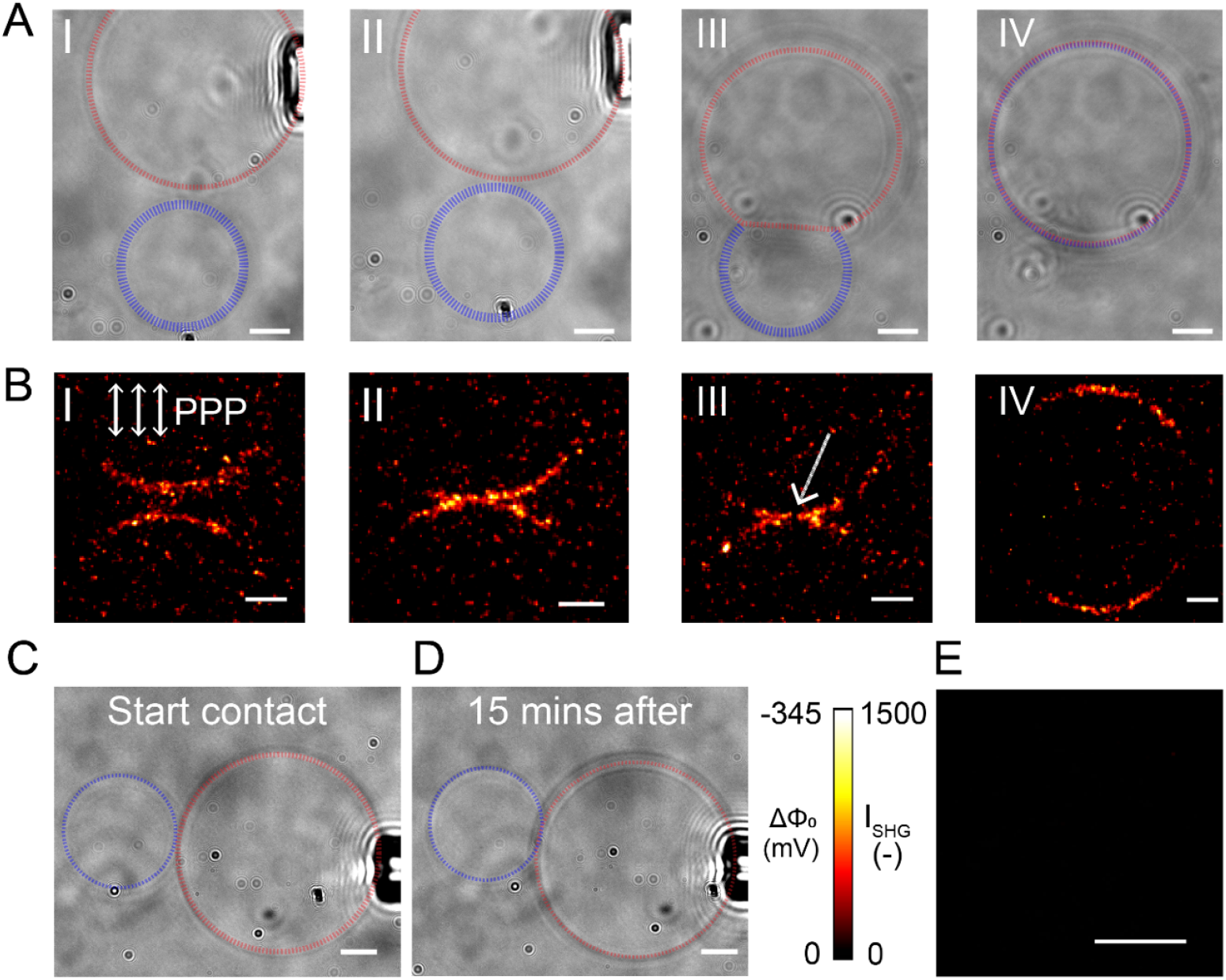
Spontaneous transmembrane potential fluctuations drive membrane fusion. **A**. White-light imaging (5 fps) of two charged GUVs composed of a 1:1 mole% ratio of DPhPC (1,2-Diphytanoyl-sn-glycero-3-phosphocholine):DPhPS (1,2-diphytanoyl-snglycero-3-phospho L-serine) that are brought into contact to fuse. The GUVs, highlighted by red and blue dashed lines, contain 15 mM sucrose internally and are suspended in an external solution of 1 mM ZnCl□ and 12 mM glucose. **B**. SH intensity and transmembrane potential (ΔΦ_0_) images of two GUVs with the same composition and media as in A undergoing fusion (calibration bar shown in E). The out-and ingoing beams were polarized perpendicular to the fusing interface (PPP polarization), as indicated by the arrows in BI. During membrane deformation (III) the SH intensity locally vanishes (white arrow), indicating that a fusion event is being initiated, and immediately following complete fusion occurs (IV). **C**,**D**. Control experiment using neutral DPhPC GUVs under identical conditions as in A, showing two GUVs initiating contact (C), and after 15 minutes of contact (D), where no fusion is observed. **E**. Potential calibration and SHG image of the neutral GUVs (as in D,E) in contact, showing no SH contrast, (no transmembrane potential). See also Video Movies S2–S3 and Fig. S9. Scale bar: 20 µm.

White-light imaging (Fig. 4A) shows a fusion sequence of two GUVs composed of DPhPS:DPhPC lipids, from approach to complete fusion. Here, one GUV is held by a micro-pipette and brought into close proximity of the other in conditions promoting transmembrane potential fluctuations (See caption and fig. S9). The dynamic progression from two shape-deformed contacted GUVs (Fig. 4A III) to a single GUV is completed within one second (Movie S2). Fig. 4B displays SH intensity and ΔΦ_0_ (transmembrane potential) images of two GUVs undergoing the same process under the same conditions as in Fig. 4A. Spontaneous ΔΦ_0_ fluctuations are visible prior to, during, and after fusion. Before contact, the two GUVs exhibit transient membrane potential fluctuations (Fig.4B I) exceeding 300 mV, sufficient for electroporation. As the two GUVs come in contact, a localized rise in ΔΦ_0_ is observed (Fig. 4B II. Measured ΔΦ_0_ histograms shown in Fig. S10). This enhancement arises from overlapping electric double layers of contact sites, increasing the probability of electroporation^8,12^ preceding the peripore-stalk–zipper pathway. Images 4A III and 4B III show a local flattening at the contact zone, relaxing membrane tension and curvature stress, as overlapping electropore formation becomes more probable (due to the larger contact areas), leading to full fusion as described earlier: The start of this process is seen as the spatial region having a vanished SH intensity (highlighted by the arrow in Fig. 4B III). GUV fusion occurs immediately afterwards, within the time resolution of the video. The fused GUV has a somewhat reduced ΔΦ_0_ compared to the initial 2 GUVs, due to the exchange of divalent ions made possible by the transient nanopore formation (Fig. 4B IV, Fig. S10). As a control, the same experiments were performed with GUVs made of net charge-neutral zwitterionic lipids kept in identical solutions, for which trans-membrane potential distributions are absent (Figs. 4C-E, where Fig. 4E shows absence of SH contrast, confirming absence of transmembrane potential). After 15 minutes of forced contact, no fusion was observed (Fig. 4C-D). These measurements demonstrate clearly that spontaneous transmembrane potentials, arising from asymmetric divalent ion distributions, are sufficient to drive membrane fusion between GUVs via electroporation with no additional fusogenic agents/electric fields, while in the absence of such potentials no such fusion occurs. These results directly validate the proposed mechanism of membrane fusion due to electroporation revealed above by the MD simulations.

## Conclusions

We describe a novel, energetically viable membrane-membrane fusion pathway driven by electroporation, where electropores, formed under a transmembrane potential, facilitate peripore stalk formation at membrane separations comparable with or greater than 1□nm—prior to the onset of hydration repulsion, and thus achievable without the need for fusogenic proteins. These peripore stalks then expand annularly, enclosing the electropores and forming full fusion pores through a “zipping” process driven by the reduction of pore edge tension. We validate our predictions by direct observation of bilayer fusion or non-fusion in the respective presence or absence of transmembrane potentials across GUV membranes. This topologically and energetically distinct pathway for membrane fusion is of particular interest because transmembrane potentials, of similar magnitude to those (ca. 0.1V/nm) leading to electropores in lipid bilayers, may occur naturally across lipid membranes of vesicles and cells, as recently demonstrated^8,12^, as well as under external application^9,34^. It furthers our understanding of membrane fusion processes, ubiquitous in areas from cell interactions to liposomic drug delivery, and highlights a further fundamental role of intrinsic membrane potential fluctuations.

## Acknowledgements

We thank Sam Safran and Pavel Jungwirth for useful discussions. Support by the European Research Council (Advanced Grant CartiLube 743016 and Synergy Grant R2-tension 951324), and the Israel Science Foundation (grant 961/2024) is gratefully acknowledged. Computational work was carried out on the Faculty of Chemistry’s high-performance computing facility CHEMFARM, which is supported in part by the Ben May Center for Chemical Theory and Computation; and supported by the WEXAC high performance computing unit and a Weizmann Institute grant for AWS cloud computing service. This work was made possible in part by the historic generosity of the Harold Perlman family.

## References

1 Bianchi, E., Doe, B., Goulding, D. & Wright, G. J. Juno is the egg Izumo receptor and is essential for mammalian fertilization. Nature 508, 483–487 (2014). 10.1038/nature13203

2 Inoue, N., Ikawa, M., Isotani, A. & Okabe, M. The immunoglobulin superfamily protein Izumo is required for sperm to fuse with eggs. Nature 434, 234–238 (2005). 10.1038/nature03362

3 Südhof, T. C. A molecular machine for neurotransmitter release: synaptotagmin and beyond. Nature Medicine 19, 1227–1231 (2013). 10.1038/nm.3338

4 Chernomordik, L. V. & Kozlov, M. M. Membrane hemifusion: Crossing a chasm in two leaps. Cell 123, 375–382 (2005). 10.1016/j.cell.2005.10.015

5 Chernomordik, L. V. & Kozlov, M. M. Mechanics of membrane fusion. Nature Structural & Molecular Biology 15, 675–683 (2008). 10.1038/nsmb.1455

6 Hed, G. & Safran, S. A. Initiation and dynamics of hemifusion in lipid bilayers. Biophysical Journal 85, 381–389 (2003). 10.1016/S0006-3495(03)74482-8

7 Jahn, R., Cafiso, D. C. & Tamm, L. K. Mechanisms of SNARE proteins in membrane fusion. Nat Rev Mol Cell Bio 25, 101–118 (2024). 10.1038/s41580-023-00668-x

8 Roesel, D., Eremchev, M., Poojari, C. S., Hub, J. S. & Roke, S. Ion-induced transient potential fluctuations facilitate pore formation and cation transport through lipid membranes. Journal of the American Chemical Society 144, 23352–23357 (2022). 10.1021/jacs.2c08543

9 Zhang, Y., Jin, D., Tivony, R., Kampf, N. & Klein, J. Cell-inspired, massive electromodulation of friction via transmembrane fields across lipid bilayers. Nature Materials 23, 1720–1727 (2024). 10.1038/s41563-024-01926-9

10 Poojari, C. S., Scherer, K. C. & Hub, J. S. Free energies of membrane stalk formation from a lipidomics perspective. Nature Communications 12, 6594 (2021). 10.1038/s41467-021-26924-2

11 Tarun, O. B., Hannesschläger, C., Pohl, P. & Roke, S. Label-free and charge-sensitive dynamic imaging of lipid membrane hydration on millisecond time scales. Proceedings of the National Academy of Sciences 115, 4081–4086 (2018). 10.1073/pnas.1719347115

12 Lee, S. et al. Dynamic second harmonic imaging of proton translocation through water needles in lipid membranes. Journal of the American Chemical Society 146, 19818–19827 (2024). 10.1021/jacs.4c02810

13 Smirnova, Y. G., Risselada, H. J. & Müller, M. Thermodynamically reversible paths of the first fusion intermediate reveal an important role for membrane anchors of fusion proteins. Proceedings of the National Academy of Sciences 116, 2571–2576 (2019). 10.1073/pnas.1818200116

14 Smirnova, Y. G., Marrink, S.-J., Lipowsky, R. & Knecht, V. Solvent-exposed tails as prestalk transition states for membrane fusion at low hydration. Journal of the American Chemical Society 132, 6710–6718 (2010). 10.1021/ja910050x

15 Kozlov, M. M. & Markin, V. S. Possible mechanism of membrane-fusion. Biofizika 28, 242–247 (1983).

16 Kozlovsky, Y. & Kozlov, M. M. Stalk model of membrane fusion: solution of energy crisis. Biophysical Journal 82, 882–895 (2002). 10.1016/S0006-3495(02)75450-7

17 Yang, L. & Huang, H. W. Observation of a membrane fusion intermediate structure. Science 297, 1877–1879 (2002). 10.1126/science.1074354

18 Fuhrmans, M., Marelli, G., Smirnova, Y. G. & Müller, M. Mechanics of membrane fusion/pore formation. Chemistry and Physics of Lipids 185, 109–128 (2015). 10.1016/j.chemphyslip.2014.07.010

19 Chanturiya, A., Chernomordik, L. V. & Zimmerberg, J. Flickering fusion pores comparable with initial exocytotic pores occur in protein-free□phospholipid□bilayers. Proceedings of the National Academy of Sciences 94, 14423–14428 (1997). 10.1073/pnas.94.26.14423

20 Golani, G. & Schwarz, U. S. High curvature promotes fusion of lipid membranes: Predictions from continuum elastic theory. Biophysical Journal 122, 1868–1882 (2023). 10.1016/j.bpj.2023.04.018

21 Rivel, T. E., Biriukov, D., Kabelka, I. & Vácha, R. Free Energy of Membrane Pore Formation and Stability from Molecular Dynamics Simulations. Journal of Chemical Information and Modeling 65, 908–920 (2025). 10.1021/acs.jcim.4c01960

22 Abiror, I. G. et al. 246 - Electric breakdown of bilayer lipid membranes I. The main experimental facts and their qualitative discussion. Bioelectrochemistry and Bioenergetics 6, 37–52 (1979). 10.1016/0302-4598(79)85005-9

23 Hsu, P. C. et al. CHARMM□GUI Martini Maker for modeling and simulation of complex bacterial membranes with lipopolysaccharides. Journal of Computational Chemistry 38, 2354–2363 (2017). 10.1002/jcc.24895

24 Jo, S., Kim, T., Iyer, V. G. & Im, W. CHARMM-GUI: A web-based graphical user interface for CHARMM. Journal of Computational Chemistry 29, 1859–1865 (2008). 10.1002/jcc.20945

25 Kotnik, T., Rems, L., Tarek, M. & Miklavi, D. Membrane Electroporation and Electropermeabilization: Mechanisms and Models. Annual Review of Biophysics 48, 63–91 (2019). 10.1146/annurev-biophys-052118-115451

26 Melcr, J., Bonhenry, D., Timr, S. t. p. n. & Jungwirth, P. Transmembrane Potential Modeling: Comparison between Methods of Constant Electric Field and Ion Imbalance. Journal of Chemical Theory and Computation 12, 2418–2425 (2016). 10.1021/acs.jctc.5b01202

27 Qi, Y. et al. CHARMM-GUI Martini Maker for Coarse-Grained Simulations with the Martini Force Field. Journal of Chemical Theory and Computation 11, 4486–4494 (2015). 10.1021/acs.jctc.5b00513

28 Ting, C. L., Awasthi, N., Müller, M. & Hub, J. S. Metastable Prepores in Tension-Free Lipid Bilayers. Physical Review Letters 120, 128103 (2018). 10.1103/PhysRevLett.120.128103

29 Vernier, P. T. et al. Nanopore Formation and Phosphatidylserine Externalization in a Phospholipid Bilayer at High Transmembrane Potential. Journal of the American Chemical Society 128, 6288–6289 (2006). 10.1021/ja0588306

30 Yarmush, M. L., Golberg, A., Serša, G., Kotnik, T. & Miklavčič, D. Electroporation-Based Technologies for Medicine: Principles, Applications, and Challenges. Annual Review of Biomedical Engineering 16, 295–320 (2014). 10.1146/annurev-bioeng-071813-104622

31 Akimov, S. A. et al. Pore formation in lipid membrane II: Energy landscape under external stress. Scientific Reports 7, 12509 (2017). 10.1038/s41598-017-12749-x

32 Jin, D. & Klein, J. The effects of splayed lipid molecules on lubrication by lipid bilayers. Lubricants 12 (2024).

33 Stevens, M. J., Hoh, J. H. & Woolf, T. B. Insights into the molecular mechanism of membrane fusion from simulation: Evidence for the association of splayed tails. Physical Review Letters 91, 188102–188102 (2003). 10.1103/PhysRevLett.91.188102

34 Haluska, C. K. et al. Time scales of membrane fusion revealed by direct imaging of vesicle fusion with high temporal resolution. Proceedings of the National Academy of Sciences 103, 15841–15846 (2006). 10.1073/pnas.0602766103

35 Kim, B. K. et al. Control of artificial membrane fusion in physiological ionic solutions beyond the limits of electroformation. Nature Communications 15, 4524–4524 (2024). 10.1038/s41467-024-48875-0

36 Saito, A. C., Ogura, T., Fujiwara, K., Murata, S. & Nomura, S. I. M. Introducing micrometer-sized artificial objects into live cells: A method for cell-giant unilamellar vesicle electrofusion. PLOS ONE 9, e106853 (2014). 10.1371/journal.pone.0106853

37 Schneck, E., Sedlmeier, F. & Netz, R. R. Hydration repulsion between biomembranes results from an interplay of dehydration and depolarization. Proceedings of the National Academy of Sciences 109, 14405–14409 (2012). 10.1073/pnas.1205811109

38 Gawrisch, K., Gaede, H. C., Mihailescu, M. & White, S. H. Hydration of POPC bilayers studied by 1H-PFG-MAS-NOESY and neutron diffraction. European Biophysics Journal 36, 281–291 (2007). 10.1007/s00249-007-0142-6

39 Murzyn, K., Róg, T., Jezierski, G., Takaoka, Y. & Pasenkiewicz-Gierula, M. Effects of phospholipid unsaturation on the membrane/water interface: A molecular simulation study. Biophysical Journal 81, 170–183 (2001). 10.1016/S0006-3495(01)75689-5

40 Hub, J. S. & Awasthi, N. Probing a Continuous Polar Defect: A Reaction Coordinate for Pore Formation in Lipid Membranes. Journal of Chemical Theory and Computation 13, 2352–2366 (2017). 10.1021/acs.jctc.7b00106

41 Dimova, R. & Marques, C. The giant vesicle book. (CRC Press, 2019).

42 Ghysels, A. et al. Permeability of membranes in the liquid ordered and liquid disordered phases. Nature Communications 10, 5616 (2019). 10.1038/s41467-019-13432-7

43 Solon, J. et al. Negative Tension Induced by Lipid Uptake. Physical Review Letters 97, 098103 (2006). 10.1103/PhysRevLett.97.098103

44 Tarun, O. B., Okur, H. I., Rangamani, P. & Roke, S. Transient domains of ordered water induced by divalent ions lead to lipid membrane curvature fluctuations. Communications Chemistry 3, 17 (2020). 10.1038/s42004-020-0263-8

